# Life as a Function: Why Transformer Architectures Struggle to Gain Genome-Level Foundational Capabilities

**DOI:** 10.1101/2025.01.13.632745

**Authors:** Hassan Hassan, Kyle Puhger, Ali Saadat, Johannes Mayer, Maximilian Sprang

**Affiliations:** DeOxy Tech, Manchester, UK; University of California, Davis, California; School of Life Sciences, Ecole Polytechnique Fédérale de Lausanne, Lausanne, Switzerland; Department of Dermatology, University Medical Center of the Johannes Gutenberg-University Mainz, Mainz, Germany, Department of Biology, Institute of Qualtitative and Computational Biology, Johannes Gutenberg University Mainz, Mainz, Germany

**Keywords:** Synthetic Biology, DNA

## Abstract

Recent advances in generative models for nucleotide sequences have shown promise, but their practical utility remains limited. In this study, we explore DNA as a complex functional representation of evolutionary processes and assess the ability of transformer-based models to capture this complexity. Through experiments with both synthetic and real DNA sequences, we demonstrate that current transformer architectures, particularly auto-regressive models relying on next-token prediction, struggle to effectively learn the underlying biological functions. Our findings suggest that these models face inherent limitations, that cannot be overcome with scale, highlighting the need for alternative approaches that incorporate evolutionary constraints and structural information. We propose potential future directions, including the integration of topological methods or the switch of modelling paradigms, to enhance the generation of genomic sequences.

## 1. Introduction

DNA/RNA molecules are the physical substrate in which life carries information. They allow for the exploration of vast combinatorial spaces and in doing so, form the backbone of evolution as a driver of change [1]. Learning how the subcomponents of these molecules are arranged to achieve specific goals (e.g. viral replication in mammalian cells) is a key problem of synthetic biology [2].

To that end, Duthie & Luque [3] have proposed a unified equation of biological evolution and population ecology. In their paper, they state, that individuals give rise to new individuals through birth such that β_i_ is the number of births attributable to individual *i*. Individuals are removed from the population through death such that δ_i_ is an indicator variable that takes a value of 1 (death of *i*) or 0 (persistence of *i*). All individuals are defined by some characteristic *z*_i_, and Δ*z*_i_ defines any change in *z*_i_ from one time step *t* to the next (*t* + 1). The total number of individuals in the population at *t* is *N*. From this foundation, we can define Ω to be a summed characteristic across *N* entities:

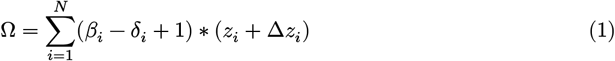

As a result of the above equation, since the rates of deaths and births are ultimately a function of the optimality of an individuals traits at a particular niche, represented by *x*, the sum of a genomes characteristics simplifies to:

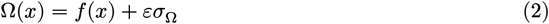

Here, *f* represents a function that selects for optimal traits, while *εσ*_Ω_ accounts for non-fatal variations. As Ω is a summation of characteristics, we expect the cumulative summation of the nucleotides of trait-relevant genes to be fairly stable at ecological niche points. This is because almost all biological traits (in the limit) are determined by nucleotides, as even epigenetic or stochastic changes are dependant on genetic capacity. Note our function here is almost always representing a stochastic (though not necessarily Brownian but usually gaussian) partial differential equation.

Focusing on DNA/RNA as the backbone of biology, many projects have aimed to learn this function using different architectures and datasets, all ultimately aiming to scale to foundational level capabilities as observed in natural language processing. Benegas et al. [4] provide a good overview of the current state-of-art for Genomic Language Models.

The key goals of this paper are to discuss two questions:

1. What is Ω?
2. What are the limits of the current paradigm with respect to learning *f*?

We hypothesize that the overall structure of Ω should be detectable and *f* learnable. The structure of Ω, in the context of genomic modeling, does not refer to 3D-chromatin structures such as topologically associated domains (TADs) as described by Rowley et al. [5], but rather to the composition of the one-dimensional symbolic representation of the genome. These include repeating elements such as tandem repeats [6], regular sequence motifs and cis-regulatory regions such as TATA boxes [7] and the differences between coding and non-coding regions [8], [9], [10], [11]. We focused our study on viral genomes, as they are small and easy to handle with low computational resources.

## 2. Ω is the output of evolution

There are a few essential points we should be aware of regarding evolution as a highly context specific, complex, adaptive framework. One, whether the framework is considered deterministic or stochastic depends on the specific particularities of the organism [12]. Two, its constraints are functional not sequential, backward not forward looking, developmental not constructive [12], [13]. There is no goal or aim to evolution. Three, much the same as selection, mutations are non-random [14]. Four, evolution’s fitness landscape is non-static [12], [15]. And finally five, it operates at the systemic level. No organism exists outside of an ecosystem [13], [15].

Taking Norwalk virus as an illustrative example, in Fig. 1 A, we can see evolutionary constraints guide sequences towards predictable patterns (particular strange attractors based on host), with deviations from those patterns correlated. This is due to the fact that each virus can only exist if it can successfully replicate, thus forcing emphasis on the genomes with the most useful traits for a particular niche (which in this case is represented by the host). Said differently, although mutations at any replication point might be Brownian and viral-cell interactions chaotic, viral genomes adapted to a host (i.e. a particular niche) approximate a temporary Lyapunov stable point to minimize system-wide energy expenditure (the system representing the viral cell populations along with the host’s cells, similar to predator-prey relationship over the longterm).

**Figure 1.**
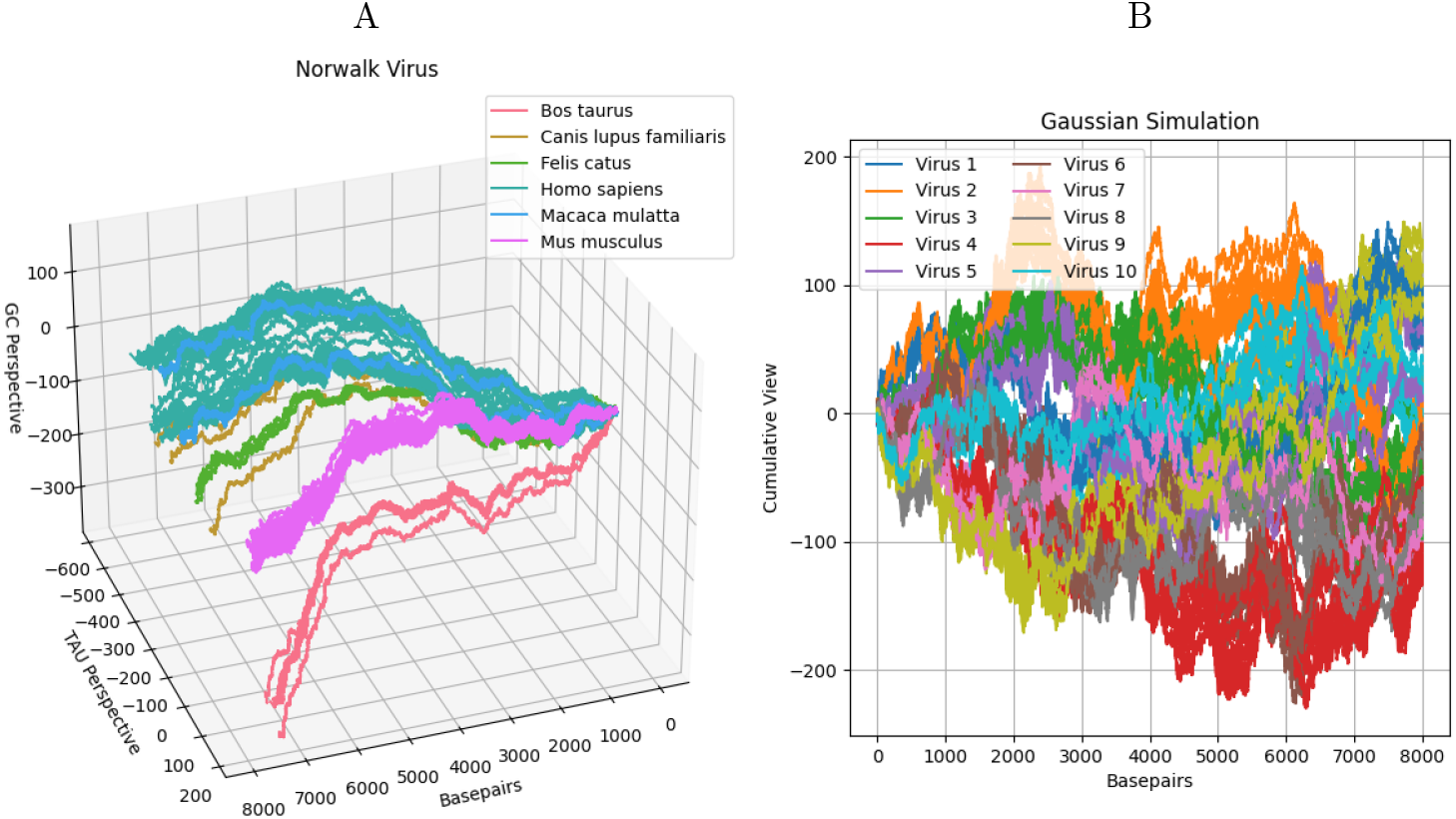
A. 3D representation of Norwalk virus samples coloured by their hosts. The basepairs axis represents the basepairs, for every basepair we move exactly one. TAU perspective represents the cumulative summation of T (0,1,1), A (0,-1,1) or U (0, 1, 1) basepairs along the y-axis. GC perspective represents the cumulative summation of G (1,0,1) and C (-1,0,1). The viruses adapted to different host species clearly occupy different regions in the overall information representation space of Norwalk virus genomes. Hosts that are closer to each other taxonomically result in overlapping regions, e.g. humans and rhesus monkeys. B. Gaussian simulation of sub-region development in the space of related viral sequences. The simulation uses 10^-5 as the mutation rate, similar to RNA viruses. x-axis represents the basepair equivalent, y-axis represents cumulative summation of selected random values, mutated by the mutation rate.

To simulate this in a Gaussian framework, we sampled 10 sequences from a random sequence matrix of the shape [samples, length, basepairs]. These 10 sequences represented successful replications within a host. Each of these sequences were then replicated 1000 times (simulating a winner takes all model) with a level of accuracy similar to viral mutation rates (10^−5^ per basepair [16]). The results of the simulation can be seen in Fig. 1 B, showing clear speciation of the initial samples, similar to what we observe in nature. What we observe here arises just from the parabolic success of a few samples from the random matrix combined with Brownian noise. As with life, the temporal and parabolic nature of replication means most sequences will contain historically induced sub-optimal components.

All together, this means that for any model to learn biological dynamics, it must learn a complex, modular function that takes into account epigenetic regulation, the regulatory networks of host cells, the constructive, interchangeable fractal nature of development [17] along with the temporal nature of evolution.

## 3. The limits of the current paradigm with respect to learning the Ω generating function

### 3.1. Transformer decoder models suffer from a power law when learning complex, modular/ compositional functions

To test the limits of transformer models in a controlled fashion, we conducted an experiment with synthetic sequences. These sequences were generated by a composite function that combined the outputs of sinusoidal terms of varying wavelengths, exponential modulation, polynomial growth, and additive noise. Samples from these functions can be found in Fig. A.4. Similar to the DNA representation above, these sequences exhibit micro- and macro-structure, although with stronger periodicity. See Appendix B.1.3 for a fuller description of the sequence-generating process.

We trained a series of Pythia-variant transformer models on these sequences, with parameter scales ranging from 1.2 million to 800 million, and evaluated their performance on out-of-distribution (OOD) sequences.

As shown in Fig. A.5 A, although model size was positively correlated with OOD performance, this correlation followed a power law, resulting in diminishing returns to scale after 300 million parameters. This is in line with the literature [18] and consistent with other experiments. Fig. A.5 B shows the training loss also stagnates (relatively) after 300 million parameters, suggesting a bound on how effectively transformer architectures can learn these functions.

#### 3.1.1 Transformer decoder models trained on real sequences minimally benefit from scale

To test whether the limits identified in our synthetic experiment extend to real DNA sequences, we repeated our experiment using the NCBI viral genomes dataset [19]. We fed viral genomes into a series of Pythia models with a context window size of 2048 basepairs and an overlap of 400 basepairs per species sample, using a simple eight-word tokenizer. See Appendix B.1.4 for more details on this process.

When trained on genetic sequences, we find results similar to those from our synthetic experiments: Transformer models need to be sufficiently large and trained on ample examples to learn from complex datasets, as shown in Fig. 2 B. However, Fig. 2 also indicates that out-of-distribution (OOD) sequences and those with few training samples (left-hand side of Fig. 2) maintain a high distance from their expected Euclidean representation. This trend persists regardless of model scale, contrasting sharply with the synthetic experiments. Therefore, our results suggest that transformers struggle to capture the underlying functions of DNA sequences, in part because their historical contingency influences how they are structured.

**Figure 2.**
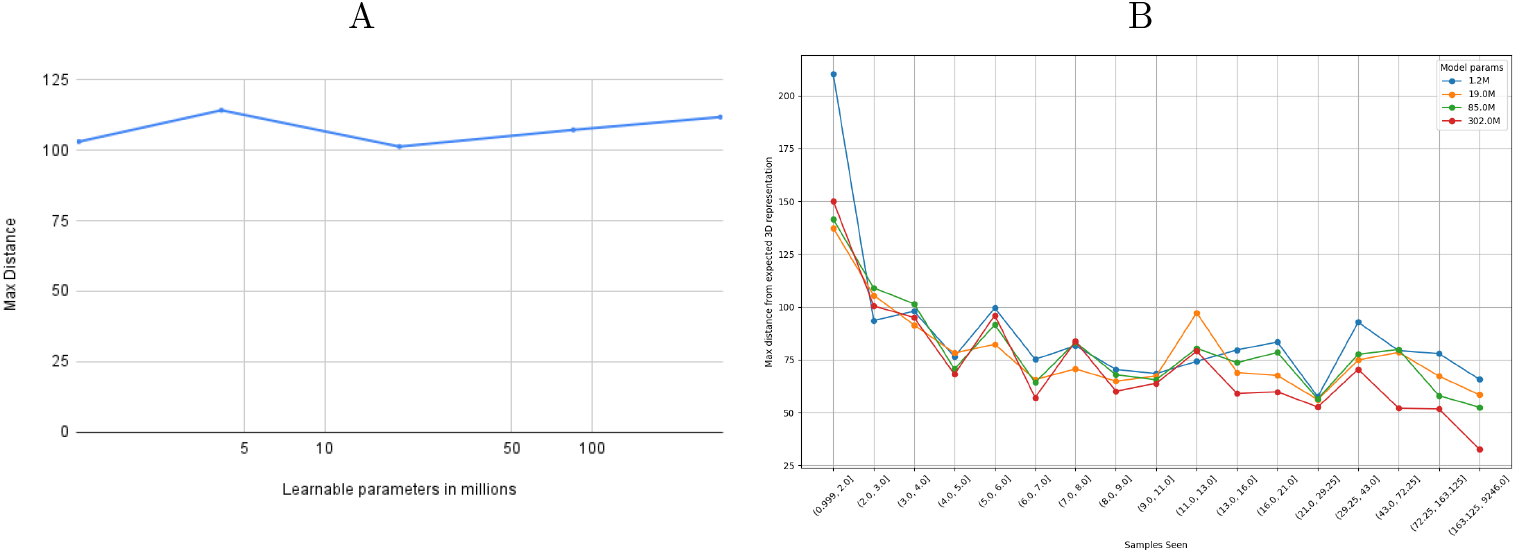
A. The first chart shows the maximum Euclidean distance (y-axis) between the expected 3D representations of OOD sequences and their generated counterparts, plotted against the logarithm of the model size (x-axis). This indicates that transformer models cannot effectively learn OOD sequences, regardless of model size. B. The second chart shows the maximum Euclidean distance between input sequences and their generated counterparts as a function of how many times each species (from which the sequence samples originate) was observed during training. Although this distance decreases as the number of examples per species increases, the overall reduction remains relatively minimal after 4 examples.

Taking a closer look at individual sequences, Fig. 3 and Fig. A.3 compare a virus with relatively few samples in the dataset (Gallivirus A) to a more abundant virus type (Enterovirus C). The true sequence is shown in black (violet in the 3D Plots), and the generated sequences are coloured by the parameter size of the generating model. In Gallivirus A (panel A), none of the models can reproduce the original sequence accurately after being prompted with the first half, although the largest model does approximate the sequence’s direction in the 3D function space. In contrast, in panel B, the two largest models learn the Enterovirus C function very well, suggesting memorisation rather than generalisation, a known issue noted in LLM research [20]. Moreover, Fig. 2 shows that OOD performance does not improve with model size, likely because OOD viral genomes originate from entirely different populations, unlike those in our synthetic experiments.

**Figure 3.**
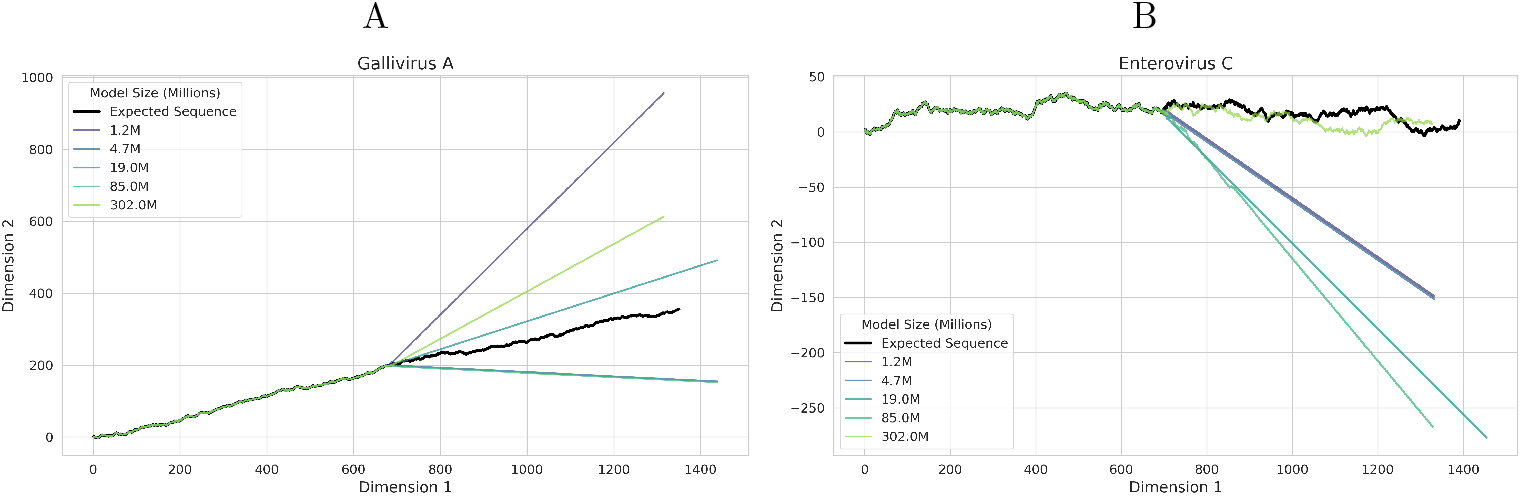
A. 2D representations of generated subsections of Gallivirus A virus sequences, coloured by model size, compared with the true sequence (in black). Each model is given half the sequence (1024 base pairs) prior to generation. No model reproduces the sequence, all are stuck in repeats. Figure axes are Cartesian coordinates of 2-dimensional vectors representing each nucleotide and then summed up to get the 2D line as by Yau et al. [21]. B. 2D representations of Enterovirus C sequences, coloured by model size, compared with the true sequence. As model size increases, output accuracy seems to improve, with the biggest model showing a generated sequence without repeats. This behavior arises from the abundance of training samples for Enterovirus C (4141) compared with Gallivirus A (15). When ample data and sufficiently large parameter sizes are available, the model memorises the species’ structure. In contrast, small sample sizes prevent memorisation and do not enable learning of underlying functions that might transfer across species. Figure axes are the same as in Fig. 1. See Fig. A.3 for a 3D representation version, were differences in direction an be observed more clearly.

### 3.2 Performance of pretrained models

Since our dataset is comparatively small, we were also interested in how larger pretrained models performed. Since Evo is one of the larger models and is an auto-regressive model, we ran it with our generation tasks.

In Figure Fig. 4 the same virus samples have been used as in Fig. 3. We observe the opposite results of our pretrained model: The Gallivirus A samples are generated quite well by Evo, although the effect, curiously, is lost when fine tuned. With Enterovirus samples, the pretrained model fails to generate the sequence well, although it does not start to repeat nucleotides immediately, the same is true for the finetuned model. This is likely due to the nature of OpenGenome, on which Evo was trained, as dataset does not contain pathogens that can infect humans (like Enterovirus), but does likely contain Gallivirus samples. These results point at memorisation again instead of generalisation.

**Figure 4.**
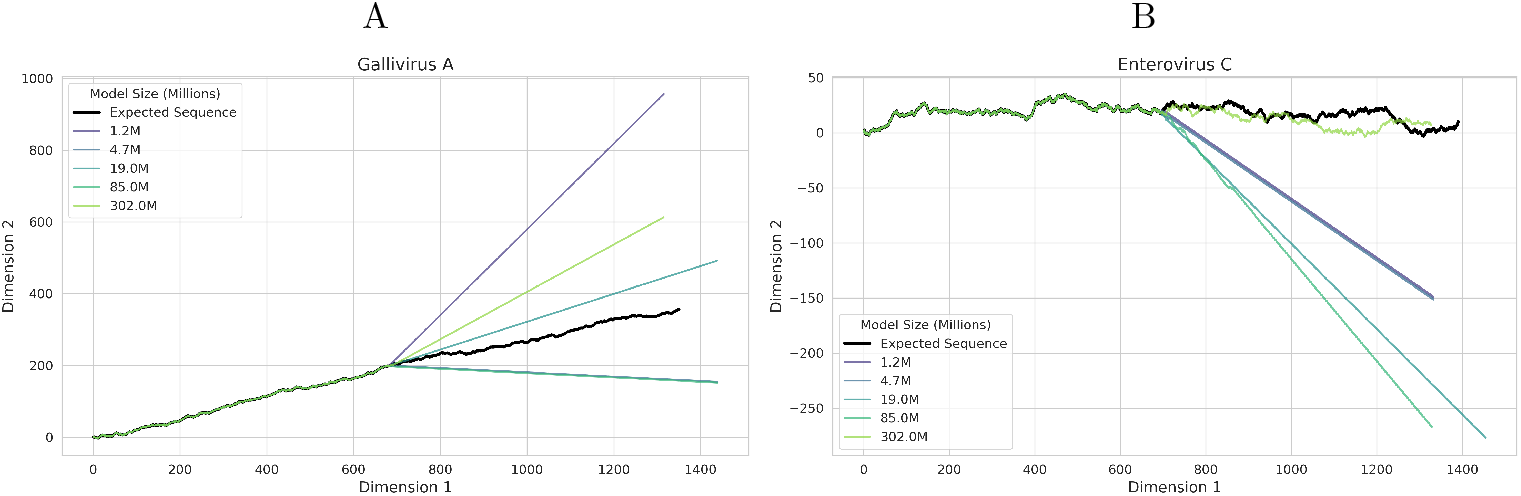
A. 2D representations of generated subsections of Gallivirus A virus sequences, coloured by model size, compared with the true sequence (in black). Each model is given half a sequence (1024 base pairs) prior to generation. No model reproduces the sequence, all are stuck in repeats. Figure axes are Cartesian coordinates of 2-dimensional vectors representing each nucleotide and then summed up to get the 2D line as by Yau et al. [21]. B. 2D representations of Enterovirus C virus sequences, coloured by model size, compared with the true sequence. As model size increases, output accuracy seems to improve, with he biggest model showing a generated sequence without repeats, but not close to the true sequence in macrostructure. This behavior arises from the abundance of training samples for Enterovirus C (4141) compared with Gallivirus A (15). When ample data and sufficiently large parameter sizes are available, the model memorizes the species’ structure. In contrast, small sample sizes prevent memorisation and do not enable learning of underlying functions that might transfer across species. Figure axes are the same as in Fig. 1. See Fig. A.3 for a 3D representation version, were differences in direction an be observed more clearly.

## 4. Discussion

In this work, we seek to provoke a discussion about the feasibility of using the currently most popular large language model (LLM) training paradigm, next-token prediction with decoder transformer models, to learn DNA sequences for use in downstream applications in the life sciences and medicine. We discuss insights from mathematical evolutionary theory that apply to the broader challenge of modeling biological dynamics, and we use them to outline what a model must learn to achieve foundational capabilities for predicting or making connections in biology. By representing sequences as a N-d line in function space, we illustrate how the functional nature of DNA becomes apparent, offering a clearer view of how viral evolution shapes the sequence function space.

Building on these insights, we use synthetic composite functions to show that transformer models struggle to learn the underlying functional structure of these sequences, exhibiting a power-law relationship for out-of-distribution (OOD) predictions. When applied to real DNA sequences, these limitations become even more pronounced: OOD samples and viral species with low abundance in the training data are reproduced poorly, resulting in large discrepancies between the original and generated sequences. These results align with findings by Vishniakov et al. [22], which also highlight a lack of foundational capabilities and broad similarity across different architectures and models.

Although transformers can be theoretically Turing-complete under certain conditions [23], our findings underscore their difficulties in learning complex functions. Augmenting transformers with “knowledge infusion” could enhance their ability to model both macro- and micro-level features of DNA. For example, topological losses, based on representations similar to those demonstrated here or in Yau et al. [21], could provide richer training signals. Similarly, persistent homology or signals analysis (e.g. amino-acid aware discrete cosine transform) based losses might enable the model to better capture large-scale structure, retain directionality, and reproduce recurring patterns during sequence generation.

Another possibility to gain foundational capabilities and generate viable DNA sequences at the genomic scale might be a switch in training paradigms/perspectives to one that addresses the nature of DNA as a function of life. There are multiple ways to model DNA as such. Firstly, we can model DNA sequences as the solution to branching random walks in a random environment as per König [24]. Due to the differing strengths of potentials within this environment, we should expect concentration of solutions to be localized islands, these islands can be viewed as our species or genera. Secondly, we can model viral DNA sequences as the phase transition between one organism and the next, gaining the capacity to build multi-scale models [25], [26]. Finally, viral genomes can be considered as sequences operating on a hierarchical topological space, with species/genera represented by topological balls, with the permeability of the topological boundaries determined by the wider energy landscape [27].

Discrete diffusion models can also be used to learn the underlying function by modelling the evolutionary genome generating path [28], [29]. As can be seen in Fig. 5, discrete diffusion models perform significantly better with OOD sampling than transformer models of comparable size when trained on the same dataset.

**Figure 5.**
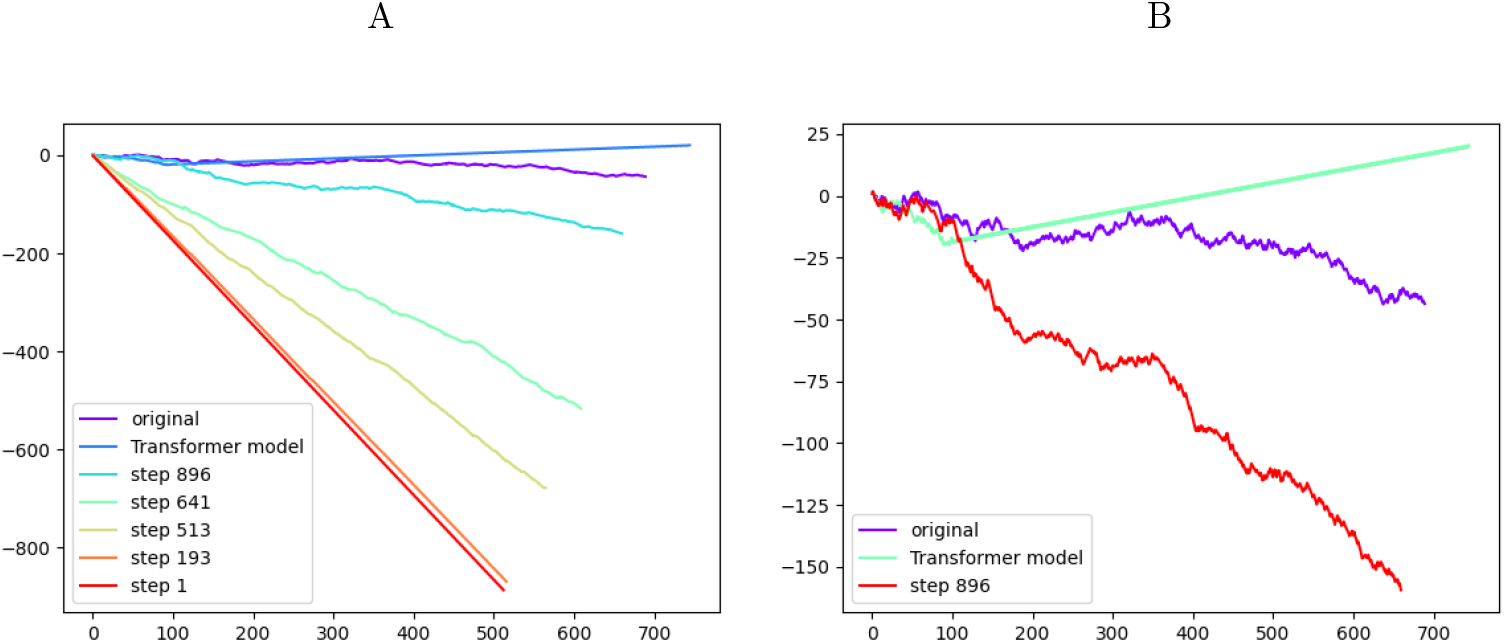
A. shows the diffusion process, by sampling from a trained model. The diffusion is guided by the first half of the sequence, comparable to the prompting of the transformer models above. The sequence is depicted by 2D-representation as per [21]. The axes are cumulative representations of the sequence similar to our 3D version, assigning a 2D-vector to each base. Colors show the different steps of the process, as well as the expected sequence (violet) and the transformer model output (blue). B. shows our selected final step for the diffusion models (red) compared to the expected sequence as well as the output of a transformer model (green) of comparable size, trained on the same data.

We can go further with flow matching models and disaggregate the problem of synthetic sequences into their natural domains (e.g. target & sequence). As can be seen in Fig. 6, flow matching models are able to generate OOD samples based just on target description.

**Figure 6.**
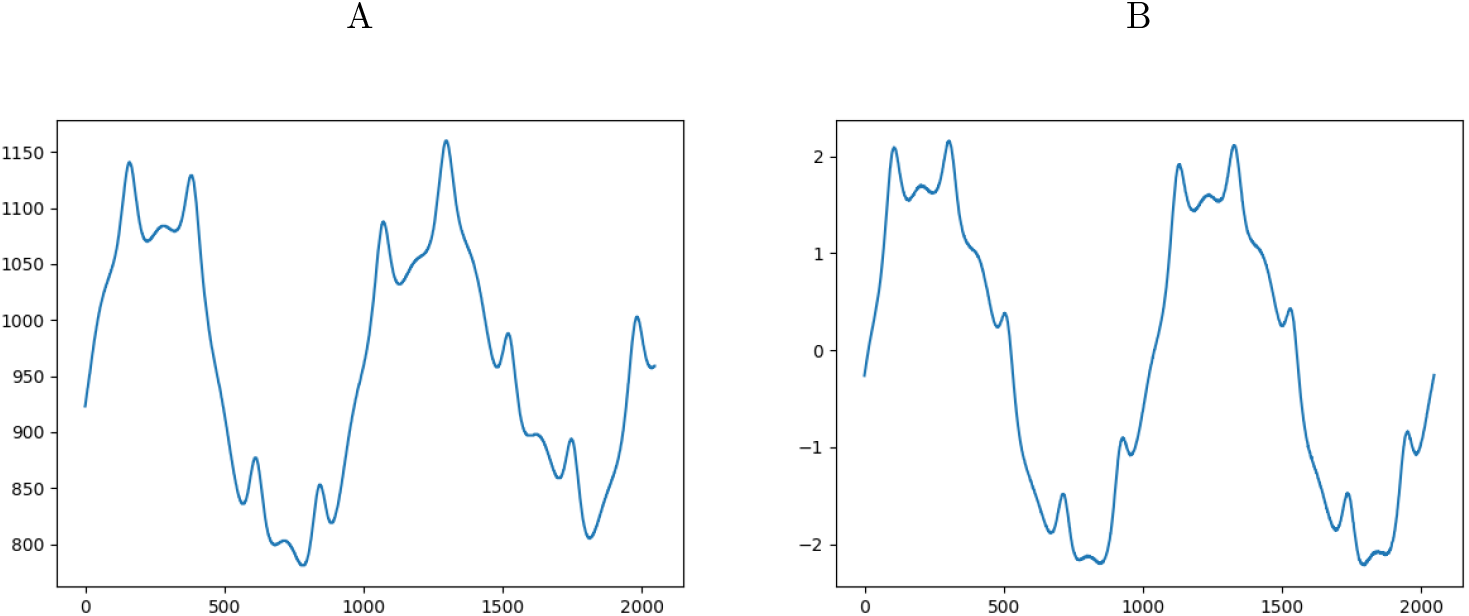
A. shows the output of a flow matching model, prompted by a target description as per Appendix B.1.3. B. shows the expected output based on the target description used to prompt the model in A.

A specific risk that should be highlighted with flow matching models, is their ease in interpolating between latents to produce potentially biologically viable results [30]. This is because they do not require their source distribution to be sampled from noise and are thus capable of learning a mapping from any distribution to another (e.g a virus hosted by one specie to another). As can be seen in Fig. 7, the model is able to learn the macro patterns associated with particular hosts. Note the chart shows GC perspective normalized and not the full view. Regardless, the dual use risk of generative models in biology must be studied and has recently started to get traction in the general discussion about the use and risk of AI [31], [32], [33].

**Figure 7.**
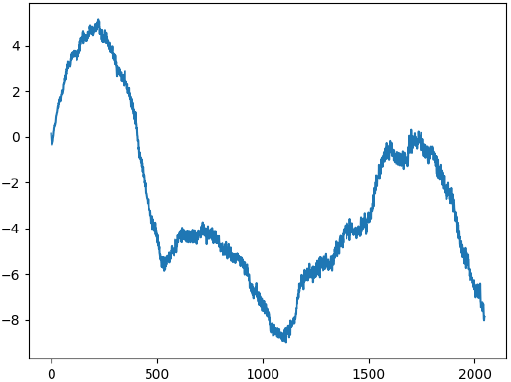
Virus interpolated from one host to another. The depicted line is the GC part of the 3D representation used in Section 3. It resembles the real distribution of GC representation of this virus species in the target host. Note we only show GC as a security measure, for this same reason, the code to recreate this experiment will not be public.

In future research we will develop flow matching models capable of generating DNA sequences at scale. Additionally, we will explore possibilities to improve the performance of next-token prediction models by designing better topological losses. During the research of these models we will aim to explore physical and in-silico risk mitigation methods, accommodating both current and potential future risks. Models of foundational capability could be applied to myriad medical use cases. Many of which will also have a strong societal impact, such as pandemic surveillance (via zoonosis prediction), synthetic bacteriophage generation for antimicrobial resistent bacteria, developing safer oncolytic phages or other biopolymers. In the light of these implications, we hope AI practitioners and stakeholders take the task of understanding the limits of foundational models seriously.

## 5 Data and Code availability

All models, code, and links to the datasets that were used will be available on GitHub at https://github.com/dna-llm/life-as-a-function. However, pre-trained models will only be available upon a reasonable request as their training data includes pathogenic viruses. All sequences were acquired off NCBI in Q2 2024.

## 6. Acknowledgements

The authors gratefully acknowledge the computing time made available to them on the highperformance computers GRETE and EMMY at GWDG at the NHR Centers NHR@Göttingen. This Center is jointly supported by the Federal Ministry of Education and Research and the state governments participating in the NHR (www.nhr-verein.de/unsere-partner). The authors thank OpenBioML (https://www.openbioml.org/) for providing a space to meet and discuss applications of machine learning in biology. Max Sprang thanks Miguel A. Andrade-Navarro for discussions and feedback and the opportunity to pursue his own projects.

## A. Appendix A

### A.1 Sequence Generation Experiments

**Figure A1:**
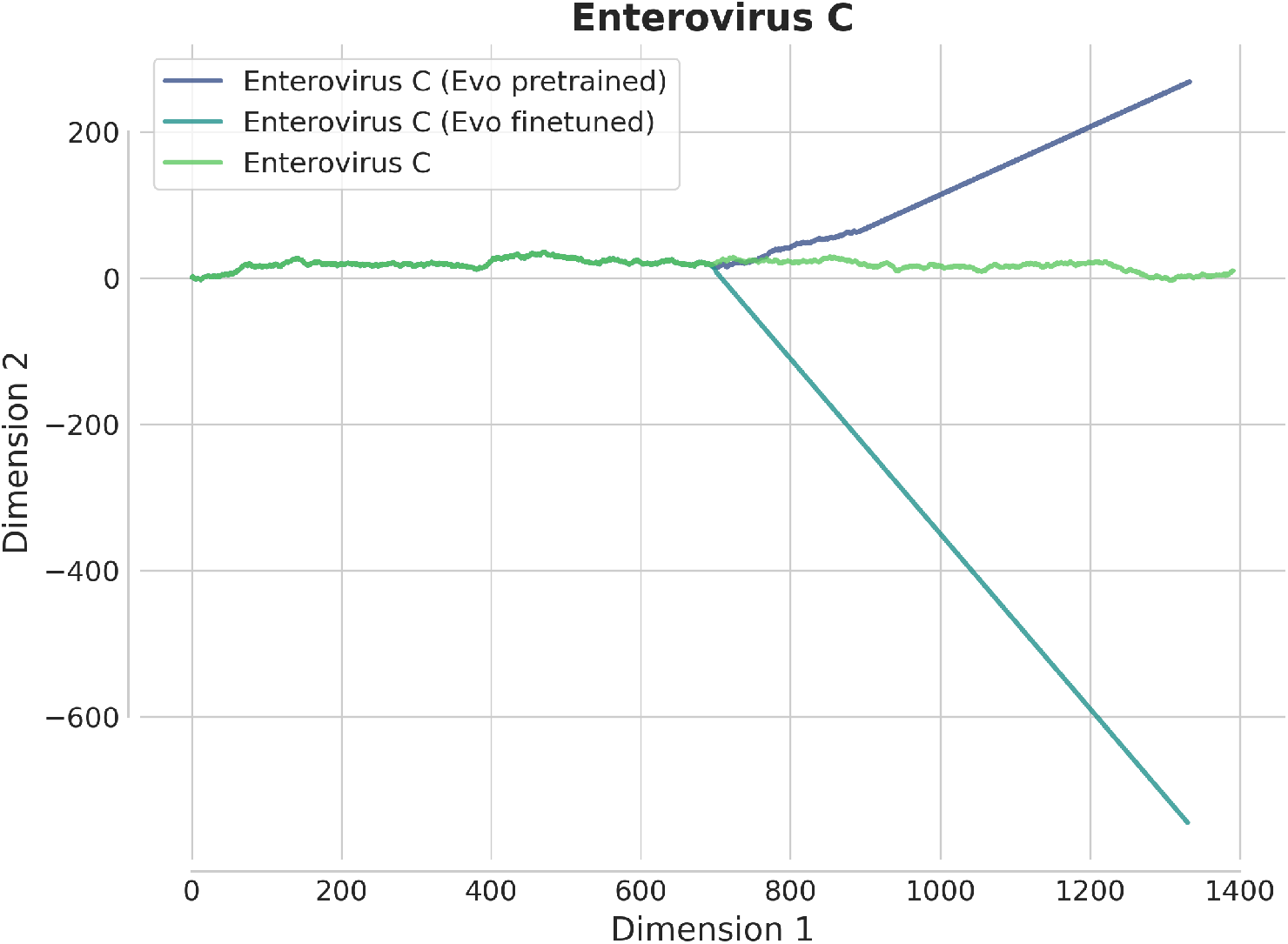
2D representations of generated subsections of Enterovirus C virus sequences, coloured by model type, compared with the true sequence (in green). Each model is given half the sequence (1024 base pairs) prior to generation. Neither model type is able to generate the required at scale. This is likely due to Enterovirus C being absent from the dataset.

**Figure A2:**
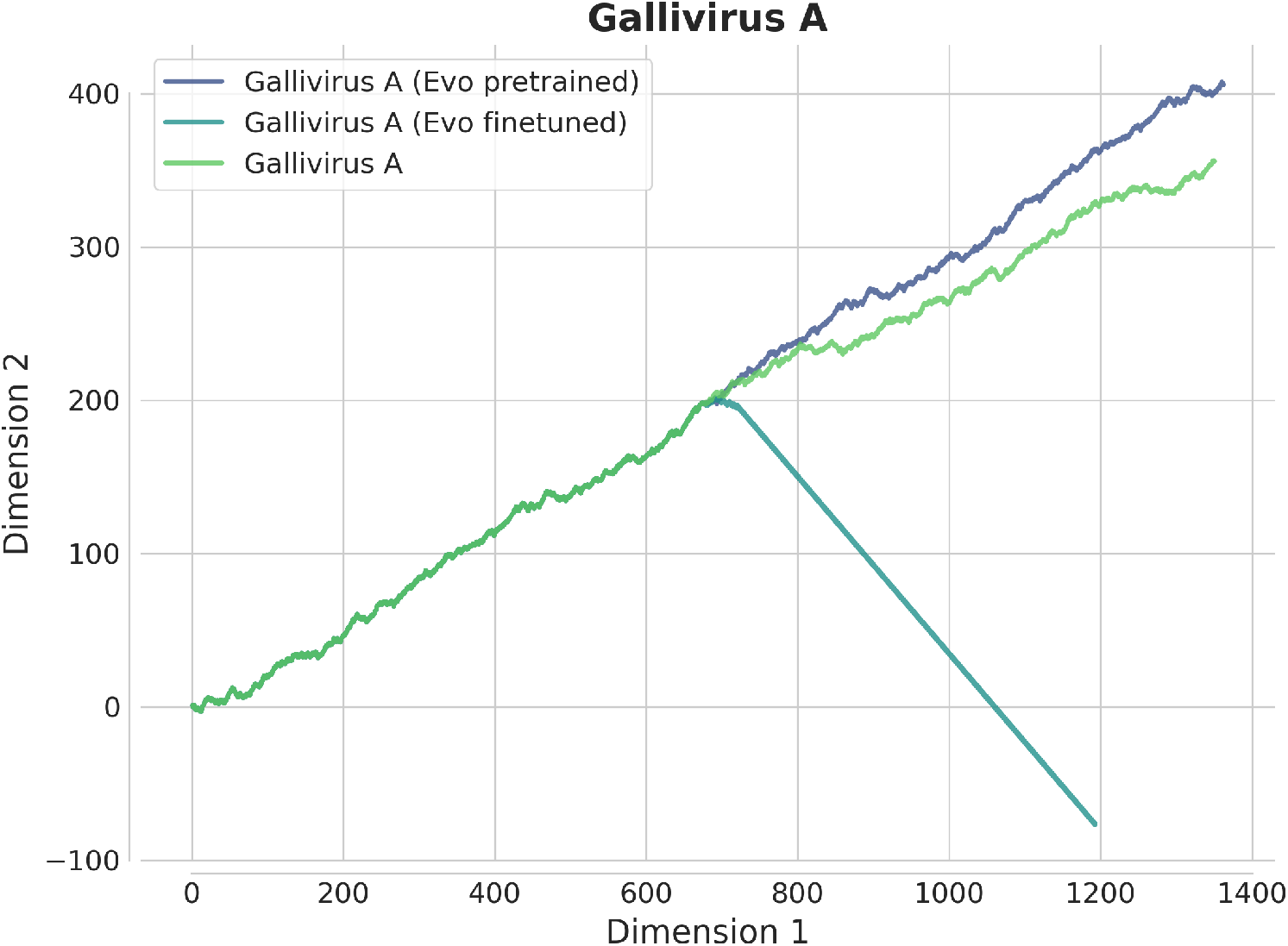
2D representations of generated subsections of Gallivirus A virus sequences, coloured by model type, compared with the true sequence (in green). Each model is given half the sequence (1024 base pairs) prior to generation. Only the Evo pretrained captures the correct representation. This is likely due to the virus being thoroughly represented data.

**Figure A3:**
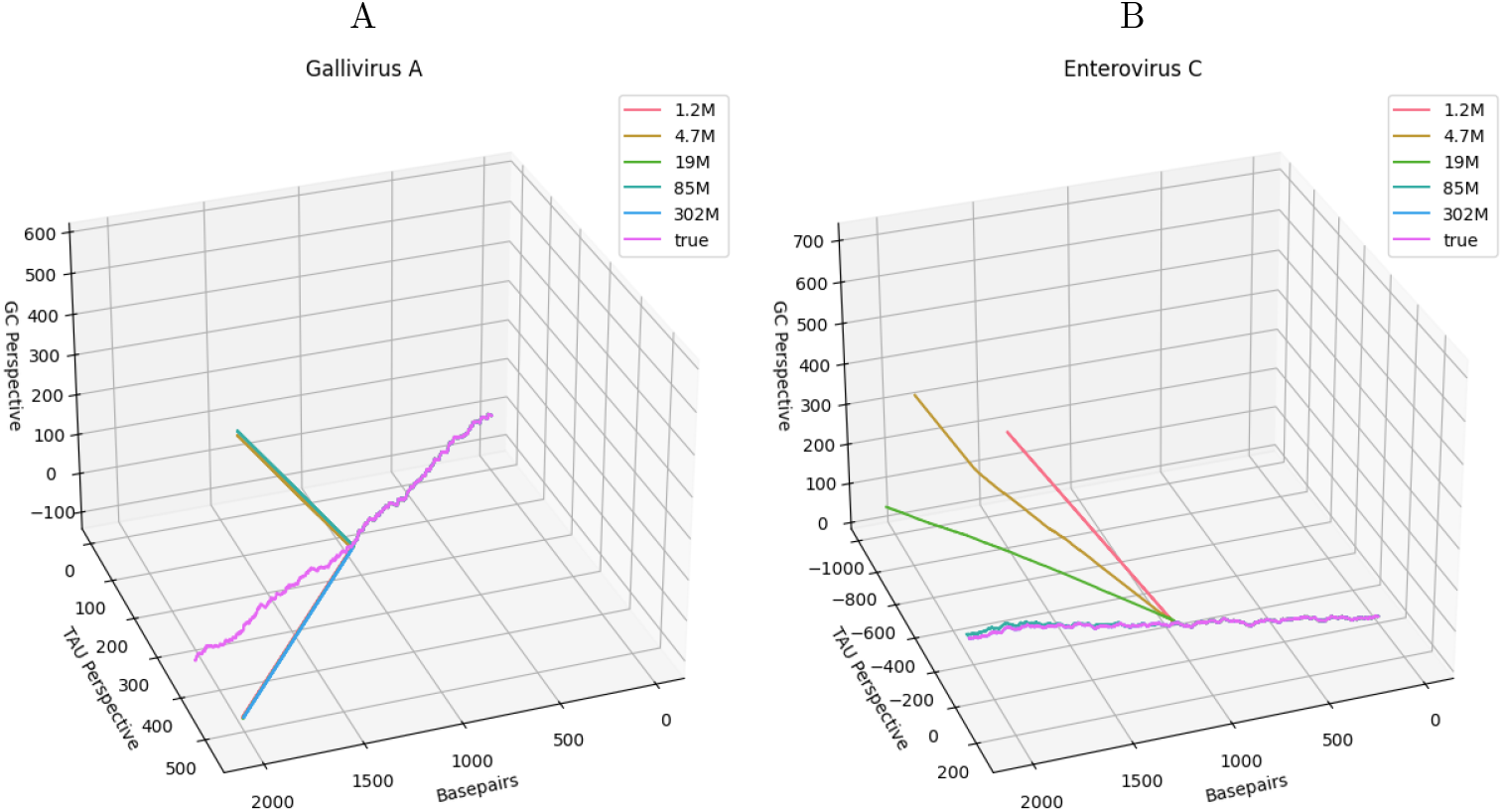
A. 3D representations of generated subsections of Gallivirus A virus sequences, coloured by model size, compared with the true sequence (in violet). Each model is given half the sequence (1024 base pairs) prior to generation. Only the 302M-parameter model captures the correct directional trend; however, none reproduces the microstructure. Figure axes are the same as in Fig. 1. B. 3D representations of Enterovirus C virus sequences, coloured by model size, compared with the true sequence. As model size increases, output accuracy improves, likely due to copying. This behavior arises from the abundance of training samples for Enterovirus C (4141) compared with Gallivirus A (15). When ample data and sufficiently large parameter sizes are available, the model memorizes the species’ structure. In contrast, small sample sizes prevent memorisation and do not enable learning of underlying functions that might transfer across species. Figure axes are the same as in Fig. 1.

### A.2. Synthetic Sequences Experiment

**Figure A4:**
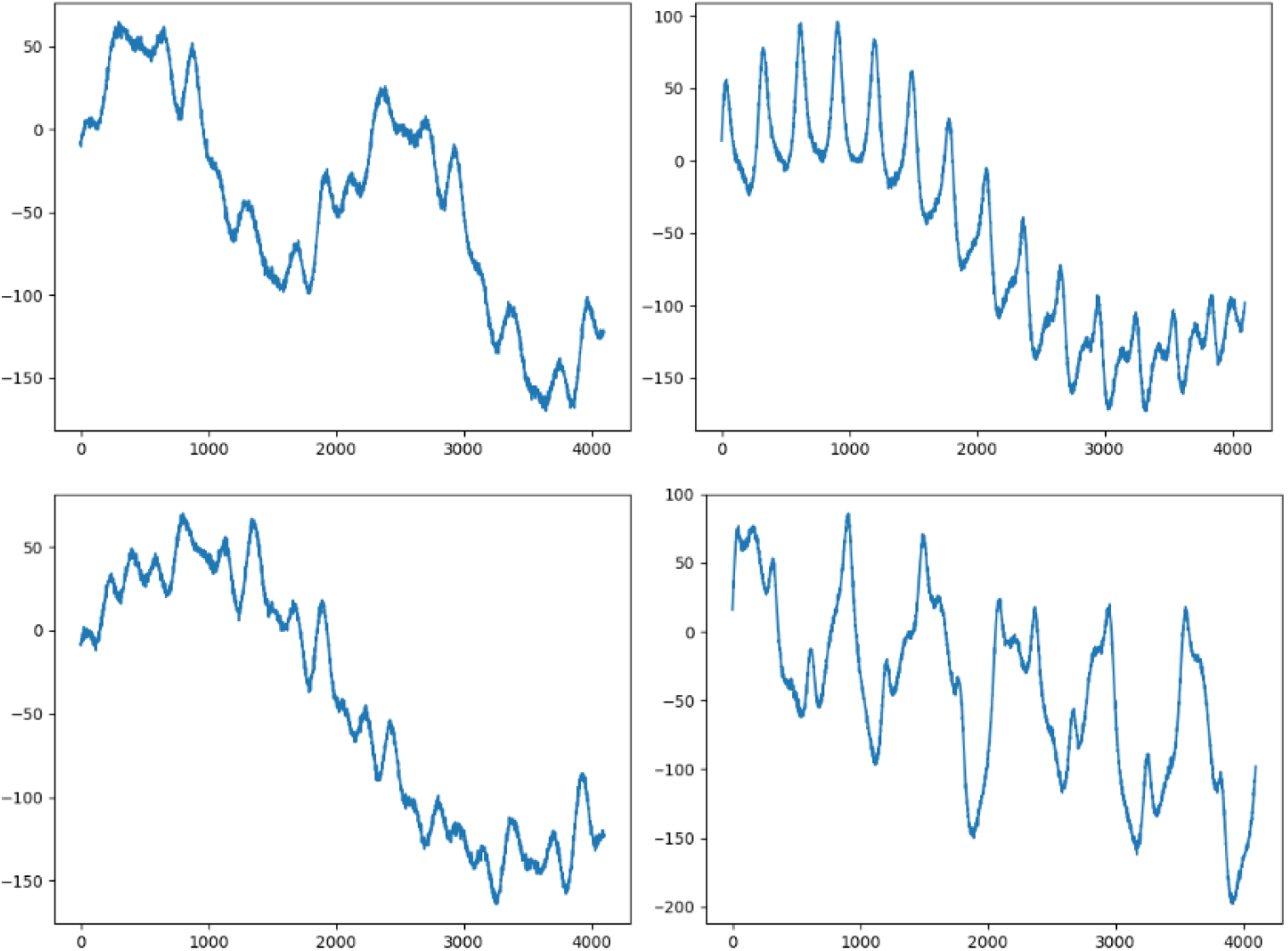
Example synthetic sequences. The x-axis represents token location and the y-axis represents the token value, which is an integer conversion of our synthetic sequence generator described in Section B.1.3.

**Figure A5:**
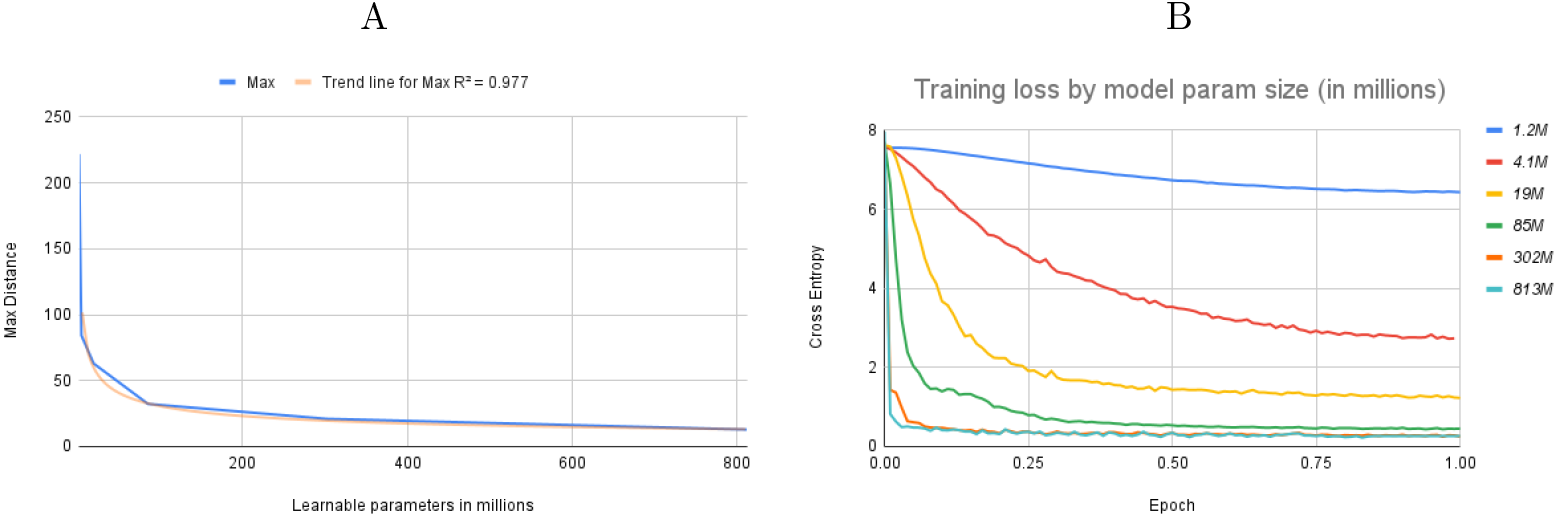
shows the maximum euclidean distance between expected 3D representations of OOD sequences and generated sequences on the y-axis against the model size on the x-axis. In contrast to DNA sequences, learning our simpler synthetic sequences improves with scale. However even then, learning is restricted to a power law, approaching negligible error decrease after 300 million parameters. B. shows cross-entropy loss across a single epoch, labelled by learnable model parameter size in millions.

## B. Appendix B

### B.1 Methods

All code, as well as additional code linked to this and potential follow up projects can be found at https://github.com/dna-llm/life-as-a-function.

#### B.1.1. Simplified 3D Representation of DNA

To investigate the spatial and energetic properties of viral DNA sequences, we utilized a dataset of viral genetic material obtained from NCBI Virus [19]. This dataset included detailed annotations of viral sequences.

To represent these sequences in a three-dimensional space, we developed a method that maps nucleotide sequences to cumulative 3D coordinates using predefined encoding schemes. The first encoding scheme, referred to as the “simplified encoding”, assigned each nucleotide a unique 3D vector based on its structural and chemical properties. The cumulative sum of these vectors across the sequence created a 3D spatial trajectory. A second encoding scheme, the “energy-based encoding”, incorporated approximations of the energetic cost of a nucleotide to capture a sequence’s energy landscape. Both mappings aimed to provide complementary insights into the spatial and energetic characteristics of a genetic sequence. We chose to go with the simplified version as we found both perspectives to be fairly similar when de-trended.

#### B.1.2. Gaussian Noise based Virus Sub-Speciation Simulation

To simulate speciation effects induced by evolution, we created 10 random matrices composed of 10,000 random numbers between 0 and 1, to symbolize initial successful viral genomes. These matrices were then copied a 1000 fold, each copy was penalized with a mutation rate, that induced a mutation at a given base pair by the chance assigned (we went with 10^−5^) mutated (i.e. a rate of every 100,000 basepairs, selected from the full genome). Note our mutations are completely random, which is different to the modern understanding of mutation [14].

#### B.1.3. Synthetic Sequence Production

To evaluate the ability of transformer models to learn complex functions, we generated synthetic sequences characterized by a combination of periodic, exponential, polynomial, and noise components. The sequence generation process was driven by a function that combined these components mathematically, incorporating sinusoidal terms with varying wavelengths, exponential modulation, polynomial growth, and additive noise. Key parameters such as amplitude, wavelength, power, and direction were systematically varied using a Cartesian product of predefined ranges. For example, wavelengths ranged from 1 to 10 units, amplitudes from 10 to 50, and amplitude ratios from 1 to 6. This systematic exploration resulted in a diverse dataset of synthetic sequences, each with a length of 4096 samples, generated using the synth_seq_gen function (find the implementation in the notebooks section of our GitHub repository). A total of 240,000 sequences were produced, and metadata for each sequence, including the parameters used, were stored for analysis. Periodically, visualizations of the generated sequences were saved to ensure quality and diversity.

The generated sequences were used to train transformer models, enabling an evaluation of their ability to approximate these synthetic functions and generalize across parameter sizes. We analyzed model performance as a function of dataset size, sequence length, and the complexity of the underlying parameters, focusing on the impact of scaling. Our results revealed that while transformers could partially learn these functions, their generalization capacity was limited, and their performance exhibited a pronounced dependence on scale. Specifically, we observed a power-law degradation in performance, highlighting intrinsic limitations in the transformer architecture for learning such structured, multi-component functions.

#### B.1.4. Experiments with Viral Genome Data

Viral DNA sequences were sourced from NCBI [19]. The dataset was filtered and only complete non-covid sequences were selected. This resulted in 210,431 samples. Each sequences was then split into 2048 basepairs with an overlap of 400 basepairs, followed by a stratified train, validation, test split.

#### B.1.5. Models

##### B.1.5.1. Transformer Architecture

We trained a Pythia transformer using the Hugging Face packages and their APIs [34].

##### B.1.5.2. Discrete Diffusion

We implemented a framework for training and evaluating a discrete flow matching model for DNA sequence modeling. The model was built on a convolutional backbone with temporal embedding layers to capture the dynamics of discrete time steps, and trained using our preprocessed DNA dataset. Key components of the pipeline included data preprocessing, training with a timestep-dependent noising schedule, and monitoring performance using FLOPs analysis.

The dataset consisted of tokenized DNA sequences from our dataset, processed into input ids for embedding. A discrete scheduler was implemented to manage the noise addition to the sequences over training steps, emphasizing harder-to-learn (more noisy) regions as training progressed. A convolutional neural network (ConvNet) was used for embedding the input sequences and modeling timestep interactions. The model incorporated layers for timestep conditioning and residual connections to capture temporal dependencies.

##### B.1.5.3. Flow Matching

To allow us to evaluate a flow-matching model’s ability to encode and generate structured sequences, we implemented a workflow combining autoencoders and a flow-based transformer model to study the representation and learning of structured synthetic sequences. Using our synthetic sequences dataset, sequences were normalized and paired with integer labels used in their generation to create a paired dataset of sequences and integers. A fully connected autoencoder, with a latent space of dimension 16, was then trained to compress and reconstruct sequences using a mean squared error loss. The encoder and decoder employed progressively narrower and wider linear layers, respectively, with dropout regularization and ReLU activations. Training spanned 15,000 steps with the Adam optimizer and noise injection for robustness.

Encoded latent representations served as input to the model, a transformer architecture configured for multi-modal learning. With a latent dimension of 16 and a transformer depth of 8, the model was trained for 100,000 steps, using gradient clipping and exponential moving average (EMA) for stability. Periodically, synthetic samples were generated by decoding latent embeddings back into sequences, and results were regularly visualized and saved. Textual labels were decoded for interpretability using a predefined mapping.

